# A northern range extension of a Canadian species of Special Concern, *Dielis pilipes* (Hymenoptera: Scoliidae), in the Okanagan Valley of British Columbia

**DOI:** 10.1101/2021.04.27.441674

**Authors:** Tyler D. Nelson, Chandra E. Moffat

## Abstract

The only known Canadian records of the yellow scarab hunter wasp, *Dielis pilipes* (Saussure) (Hymenoptera: Scoliidae), are from the southern Okanagan and Similkameen valleys of British Columbia. We report a 25-km northern range extension of the species into the Ponderosa Pine Biogeoclimatic Ecosystem Classification zone, collected in an unmanaged agricultural field in Summerland, British Columbia. This finding is of conservation importance and has implications for natural biological control of ten-lined June beetles, *Polyphylla decemlineata* (Say) and *P. crinita* LeConte (Coleoptera: Scarabaeidae), incidental agricultural pests in the Okanagan.

---

In Canada, the yellow scarab hunter wasp, *Dielis pilipes* (Saussure) (Hymenoptera: Scoliidae) is known only from the southern Okanagan and Similkameen valleys of British Columbia, where it is closely associated with antelope brush, *Purshia tridentata* (Pursh.) de Candolle (Rosaceae), and sagebrush, *Artemisia* spp. Linnaeus (Asteraceae), ecological communities below 600 m elevation (COSEWIC 2018). The Committee on the Status of Endangered Wildlife in Canada (COSEWIC) recently designated *D. pilipes* as a species of Special Concern based on loss, degradation, and fragmentation of these habitats, in addition to pesticide use in adjacent agriculture land. Their report summarises the 68 known Canadian records of the species across 14 sites (Fig. 1; J. Heron and C. Sheffield 2020, personal data), almost all of which are in the 40-km area between Osoyoos and northern Okanagan Falls in the Okanagan Valley in the Bunchgrass Biogeoclimatic Ecosystem Classification zone (Meidinger and Pojar 1991; COSEWIC 2018; Mackenzie and Meidinger 2018). Here, we report five new *D. pilipes* records collected in Summerland, British Columbia extending its confirmed range 25 km north of the COSEWIC records into the Ponderosa Pine Biogeoclimatic Ecosystem Classification (BEC) zone.

**Fig. 1.**
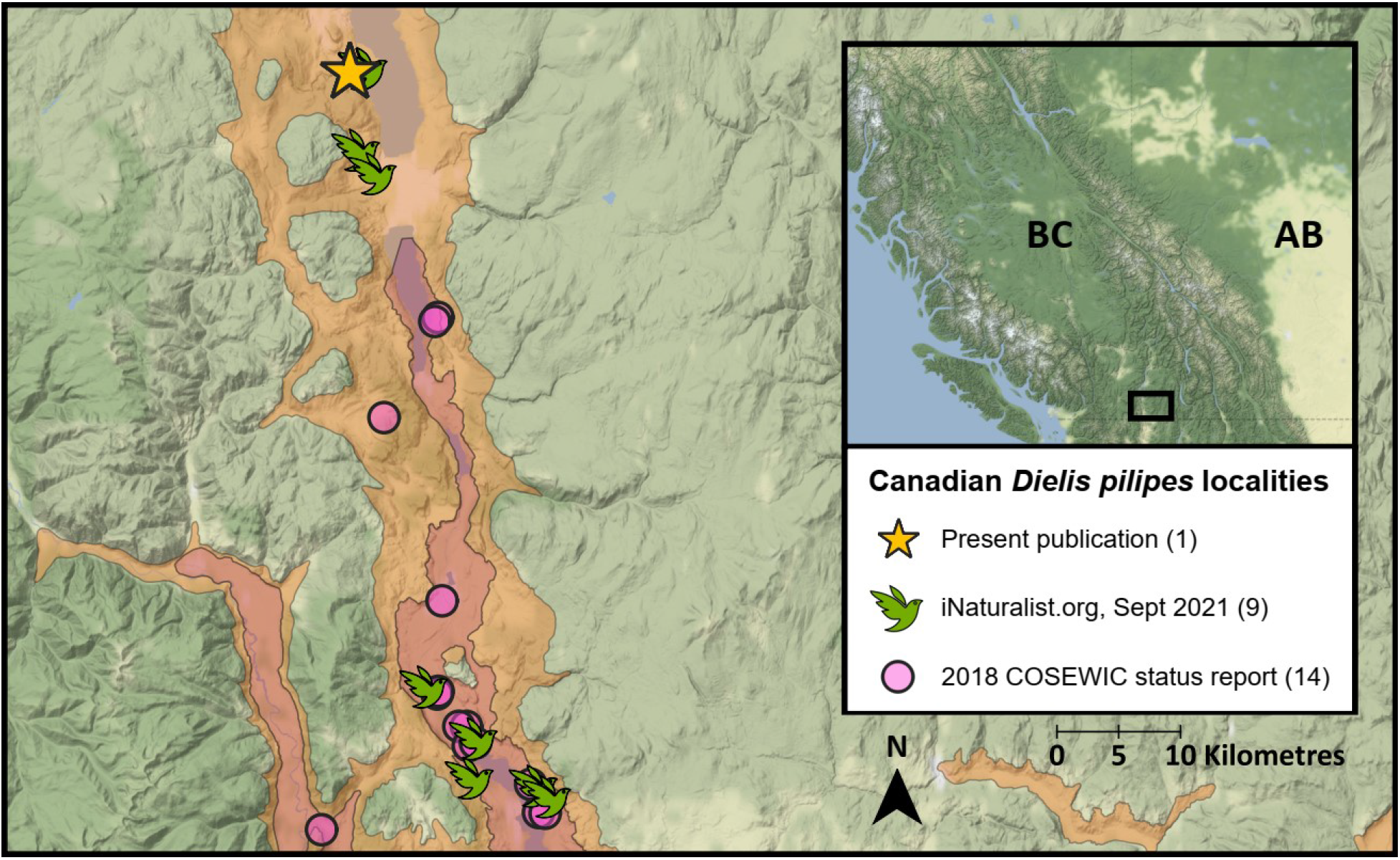
All recorded *Dielis pilipes* (Saussure) occurrences in Canada and their sources. Orange map area indicates Ponderosa Pine Biogeoclimatic Ecosystem Classification zone; red indicates Bunchgrass zone. Number of localities from each source are in parentheses. COSEWIC occurrence records provided by J. Heron and C. Sheffield (unpublished data). Figure prepared in QGIS, version 3.6.0, and base map obtained from https://github.com/stamen/terrain-classic.

Adult *D. pilipes* are large, black wasps with yellow bands on their first 3–5 abdominal tergites (MacKay 1987). In British Columbia, the species has been observed nectaring on showy milkweed (*Asclepias speciosa* Torrey (Apocynaceae)), alfalfa (*Medicago sativa* Linnaeus (Fabaceae)), and white sweet-clover (*Melilotus albus* Medikus (Fabaceae)), but other plants that flower during their flight period are probable nectar sources (COSEWIC 2018). Adult female scoliids lay single eggs on larvae of scarab beetles (Coleoptera: Scarabaeidae), and the larvae develop externally on their host (Krombein 1979; O’Neill 2001). Although reproduction of *D. pilipes* has not been observed in Canada, the species will parasitise the ten-lined June beetle, *Polyphylla decemlineta* (Say) (Coleoptera: Scarabaeidae), in the United States. *Polyphylla crinita* LeConte (Coleoptera: Scarabaeidae) is a probable host of the wasp, although this remains undocumented (COSEWIC 2018). These two scarabs are found throughout British Columbia (Bousquet *et al.* 2013), far beyond the known range of *D. pilipes*, but it is unclear what factors restrict the wasps to the Okanagan and Similkameen valleys (COSEWIC 2018). The scarabs can cause considerable economic damage in agricultural landscapes across western North America (Downes and Andison 1941; Van Steenwyk and Rough 1989).

We collected five *D. pilipes* that were nectaring on Brassicaceae flowers in a flat, disturbed, and unmanaged agricultural field on the grounds of Agriculture and Agri-Food Canada’s Summerland Research and Development Centre in Summerland, British Columbia, Canada (49° 33’ 45.58” N, 119° 39’ 07.48” W) between 19 June and 8 July 2020 (Fig. 1). The field is a remnant patch of grassland shrub–steppe located in the Ponderosa Pine BEC zone, subzone variant xh1 (Okanagan Very Dry Hot; Meidinger and Pojar 1991; Mackenzie and Meidinger 2018). The lead author (TDN) visited the site at least once per week between 28 May and 10 July 2020, after which visits occurred every second or third week until 8 October 2020 (Table 1). All visits were between four and seven hours in length, typically beginning around 10:00 a.m. Dominant forb species in the site were alfalfa, stork’s-bill (*Erodium cicutarium* (Linnaeus) L’Héritier de Brutelle ex Aiton (Geraniaceae)), Russian thistle (*Kali tragus* (Linnaeus) Scopoli (Amaranthaceae)), and baby’s breath (*Gypsophila paniculata* Linnaeus (Caryophyllaceae)). All *D. pilipes* were collected haphazardly by aerial net while measuring plant growth in an experimental plot; TDN collected one on 19 June, one on 3 July, and three on 8 July 2020. We identified each as female *D. pilipes* (Fig. 2), using the key in MacKay (1987). TDN also collected one adult female *Polyphylla* from the site on 24 July 2020. In addition to these collections, TDN observed 5–10 *D. pilipes* nectaring nearby the site (49° 33’ 46.65” N, 119° 39’ 00.23” W) on one occasion in early July 2020.

**Table 1.**
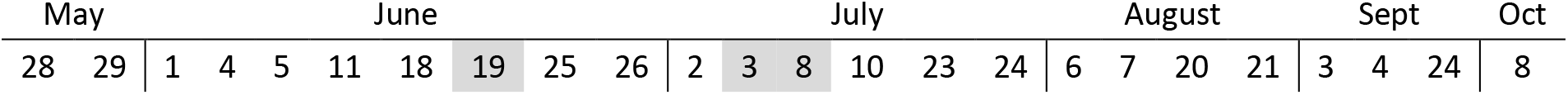
Calendar days of 2020 when the authors were present in the *Dielis pilipes* (Saussure) site in Summerland, British Columbia (49° 33’ 45.58” N, 119° 39’ 7.48” W). Grey boxes denote specimen collection events.

**Fig. 2.**
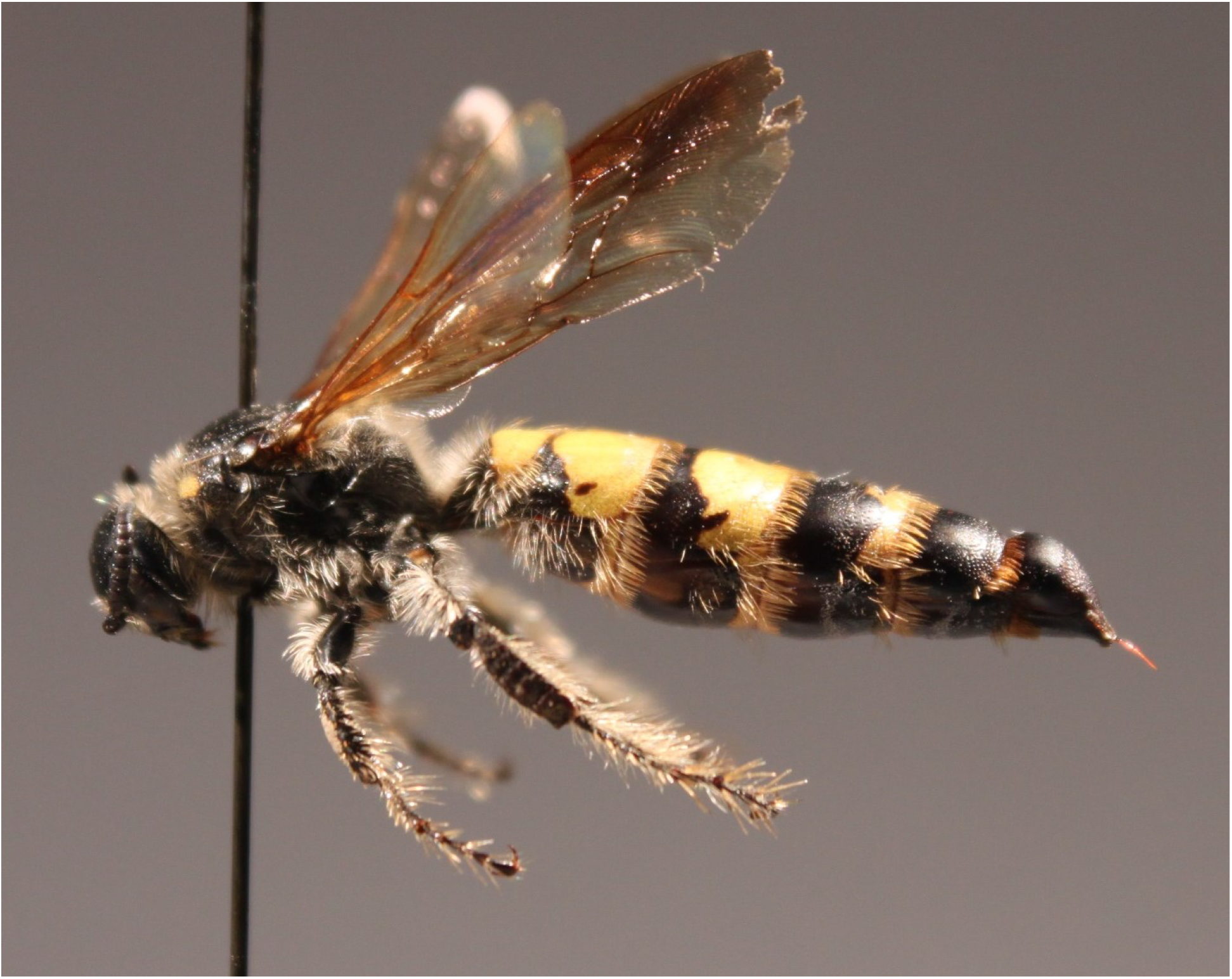
A female *Dielis pilipes* (Saussure) collected in Summerland, British Columbia.

We report a northern range extension of *D. pilipes* into the Ponderosa Pine BEC zone. Before this report, only one *D. pilipes* individual had been recorded in this zone; however, the exact location of this record is unclear (Fig 1; J. Heron and C. Sheffield 2020, unpublished data). This range extension is supported by our five specimens from 2020, four observations in 2021, as well as additional research-grade observations uploaded to iNaturalist.org (iNaturalist.org 2021). We observed adult flight at the northernmost extent of its range to be between 19 June and 8 July 2020. These findings provide additional evidence of its scarcity in British Columbia (Table 1; see COSEWIC 2018). We suspect that the species has only recently established in the Summerland area; it was not detected in any conservation assessment survey north of Penticton, British Columbia (COSEWIC 2018, Fig. 8), and no specimens are housed in the Summerland Research and Development Centre’s insect collection despite decades of on-site collecting by federal research staff.

The detection of *D. pilipes* in Summerland could have implications for pest management in the central Okanagan, as these wasps may be impacted by pesticide sprays in local orchards (COSEWIC 2018). In addition, the species is a natural biological control agent of *Polyphylla* beetles and therefore are valuable for orchard managers who have few other options to manage belowground beetle larvae (COSEWIC 2018). We recommend that future survey work for *D. pilipes* include sites in or nearby the grounds of the Summerland Research and Development Centre to determine future locality occupancy. In addition, we recommend surveys further north in the Okanagan Valley because we expect to see the species continue its expansion northwards into the Ponderosa Pine BEC zone. Vouchers of *D. pilipes* have been deposited in the centre’s collection and at the Royal British Columbia Museum in Victoria, British Columbia.

## Acknowledgements

The authors thank Jennifer Heron, Cory Sheffield, and Richard Cannings for providing Canadian *Dielis pilipes* records. They also thank the AAFC Summerland Research and Development Centre field staff and David Ensing for their cooperation. They again thank David Ensing and two anonymous reviewers for their helpful comments on the manuscript. The Biodiversity Heritage Library (https://www.biodiversitylibrary.org/) supported this study by hosting historical publications, which were difficult to access otherwise.

